# The alkaliphilic side of *Staphylococcus aureus*

**DOI:** 10.1101/735191

**Authors:** Manisha Vaish, Amyeo Jereen, Amall Ali, Terry Ann Krulwich

**Author notes:** To whom correspondence may be addressed. Address: Department of Pharmacological Sciences, Icahn School of Medicine at Mount Sinai, Box 1603, 1 G. Levy Place, New York, NY 10029. Department of Pharmacological Sciences, Icahn School of Medicine at Mount Sinai, New York, NY 10029. A.J.: 12901 Bruce B Downs Blvd, Tampa, FL 33612. A.A.: 98 Morningside avenue, Apt 54, New York, NY 10027. TAK: 1160 Park Avenue, New York, NY 10128-1212. **Email (E) and phone (ph) numbers of all co-authors** Jereen: E; ph: none Ali: E; Krulwich: E.

## Abstract

The genome of *Staphylococcus aureus* has eight structurally distinct <u>c</u>ation/<u>p</u>roton <u>a</u>ntiporters (CPA) that play significant roles in maintaining cytoplasmic pH and ions in extreme conditions. These antiporters enable *S. aureus* to persist under conditions that are favorable to the bacterium but unfavorable to animal host including humans. In this study, we report physiological roles and catalytic properties of NhaC (NhaC1, NhaC2 and NhaC3), CPA1 (CPA1-1 and CPA1-2) and CPA2 family antiporters and how these antiporters crosstalk with Mnh1, a CPA3 family antiporter, recently shown to play important roles in virulence and pH tolerance. Catalytic properties of antiporters were determined by Na^+^/H^+^ and K^+^/H^+^ antiport assays using everted membrane vesicles of a CPA-deficient *E. coli* KNabC host. NhaC and CPA1 candidates exhibited Na^+^/H^+^ and K^+^/H^+^ antiporter activity in the pH range between pH 7 to 9.5 but did not show significant role in halotolerance and osmotolerance alone. Interestingly, NhaC3 exhibited significant antiporter activity at alkaline pH and play major roles in pH and salt tolerance. CPA2 neither exhibited Na^+^or K^+^/H^+^ exchange nor showed any active role in pH and salt tolerance. Double deletion of *mnhA1* with *nhaC1, nhaC3, cpa1-1 or cpa1-2* respectively, made *S. aureus* severely sensitive at pH 7.5 under stress conditions indicating synergistic relationship of Mnh1 with these antiporters. The functional loss study of these antiporters in *in-vivo* mouse infection model, *nhaC3* deletion showed significant loss of *S. aureus* virulence. Altogether, the current study indicates NhaC3 as a potential target against *S. aureus* virulence under extreme pH and salt conditions.

**Importance:** In this study, we established catalytic properties and physiological roles of *S. aureus* NhaC, CPA1 and CPA2 family antiporters and their importance under salt and alkaline stress conditions. Except CPA2, all five antiporters of both families were active for Na^+^/H^+^ and K^+^/H^+^ exchange. CPA1-1 showed significant role in pH homeostasis at pH 7.5 whereas CPA1-2 and NhaCs were major contributors to halotolerance and osmotolerance at alkaline pH. The severity of growth deficit in double knockouts of *mnhA1* with each of *nhaC1, nhaC2, nhaC3, cpa1-1* or *cpa1-2* establishes their synergistic relationship in regulating pH and salt homeostasis. Deletion of *cpa1-1, cpa1-2* and *nhaC1, nhaC2*, and *nhaC3* were assessed in mice model and NhaC3 was shown to play a major role in *S. aureus* virulence.

## Introduction

*S. aureus* has a robust defense mechanism that ensures its survival in extreme stress conditions within a host as well as in the environment. It can successfully colonize in stomach, small intestine, skin, and external nares where extreme pH and salt conditions exist. Adaptation of *S. aureus* in diverse niches leads to infection ranging from minor skin infections like abscesses and boils to life threatening diseases like osteomylitis, endocarditis and food poisoning etc. (1).

The survival of *S. aureus* in high salt and alkaline environment highly depends on the active secondary transporters on the cell membrane. This group of secondary transporters belongs to monovalent cation/proton antiporter (CPA) family. They take up external protons utilizing inward proton gradients generated by respiration or other specific proton pumps. Concomitantly, these antiporters catalyze efflux of cytoplasmic cations that are potentially toxic, e.g. Na^+^, Li^+^ or excess K^+^ (2). Such antiporters were first observed in alkaliphilic *Bacillus firmus* OF4 (later renamed *B. pseudofirmus* OF4), were named <u>N</u>a^+^/<u>H</u>^+^ <u>a</u>ntiporters type <u>C</u> (NhaC) (3) and now have 4 numbered sub-sets, e.g. containing antiporter variants such as the H^+^-malate/Na^+^-lactate antiporters or new species (4). These antiporters play major role in proton retention and in maintaining intra-cellular pH homeostasis at highly alkaline conditions.

Extremophiles inhabit extreme alkaline environments where pH > 12 such as soda lakes and in industrial settings such as indigo dye plants, sewage plants, and underground water etc. (5, 6). These extreme alkaliphiles use many of the same strategies observed in neutralophiles, further adapting them to respond to more extreme challenges. Typically, proteins involved in the pH homeostasis mechanisms of extremophiles are constitutively expressed, so that these bacteria are prepared for sudden shifts to the extreme end of the pH range. Similarly, neutralophiles maintain substantially more acidic cytoplasmic pH than the external pH at the higher end of their pH range. Bacteria have additional strategies for surviving without growth during periods of exposure to pH values that are outside their growth range. Survival without growth is assessed by the resumption of growth on return of the bacteria to a permissive pH (i.e., a near-neutral pH for neutralophiles). For example, *S. aureus*, a neutralophile that can grow at external pH values of 5.5 to 9.5 but generally maintains its cytoplasmic pH between 7.4-7.7 (7). Enteric bacteria such as *Escherichia coli* and *Salmonella* spp. survive passage through the stomach but do not grow in that niche (2, 8) and *E. coli* survives exposure to alkaline sea water but does not grow (9).

Survival and growth under acidic or alkaline stress involves changes in the cell structure, metabolism, and transport patterns. Cell membrane transporters play a major role in maintaining homeostasis within the cell during pH shift in the environment. The groups of secondary transporters, mainly Na^+^/H^+^ antiporters, have been found in many alkaliphiles and pathogenic bacteria important for their physiology, ecology and pathogenesis. For alkaliphiles, Na^+^/H^+^ antiporters are not essential to maintain alkaliphilic nature but play an important role in regulating cytoplasmic pH at higher environment pH (10, 11). In some of the major pathogens, cation/proton antiporters have been linked with virulence. Transporters in *E. coli* B2 strain have an important role in extra intestinal virulence without strong effect on commensalism and these could serve as drug target against *E. coli* infections (12, 13). The NhaA-type Na^+^/H^+^ antiporter of the Bubonic plague pathogen, *Yersinia pestis*, has been shown to be essential for its virulence (14). The NhaA-type Na^+^/H^+^ antiporter of *Vibrio cholera* has been shown to be important for the viability of the pathogen in its Na^+^ rich biotope (15). NhaP2, a K^+^/Na^+^/H^+^ antiporter of *Vibrio cholera*, has been shown to be important for its adaptation to acidic environment (16, 17). Recently, we found that a multi-drug resistant pathogen *S. aureus* seven-subunit cation/proton antiporter Mnh1 of CPA3 family plays important role in diminishing sodium toxicity and maintaining pH homeostasis at neutral pH. We found Mnh1 to be essential for virulence and pathogenesis of *S. aureus* in the *in-vivo* mice infection model, and therefore it could be a potential therapeutic target (18).

In this study, we are reporting NhaC, CPA1 and CPA2 family antiporters with their catalytic properties and roles in physiology, ecology, and pathogenesis of *S. aureus* for the first time.

## Results

### Characterization of catalytic properties of NhaC, CPA1 and CPA2 antiporters

Catalytic properties of NhaC1, NhaC2, NhaC3, CPA1-1, CPA1-2 and CPA2 were determined by the antiporter assay. These six *S. aureus* Newman antiporters were cloned into inducible pBAD vector and transformed into the cation/proton antiporter deficient *E. coli* KNabC strain, keeping empty pBAD vector as control. Everted membrane vesicles were prepared using very high pressure and antiport assays were performed at pH range 7.0-9.5. NhaC candidates exhibited catalytic activities for Na^+^/H^+^ and K^+^/H^+^ exchange at neutral to alkaline pH ranging 7.0-9.5. They exhibited modest activities in the pH range between 7.0 to 8.0 in presence of high salt concentration (1 M NaCl or 1M KCl). Increasing pH accelerated antiport activity and the activity was optimum at pH 9.5 (Fig 1). At this pH, NhaC3 efficiently showed Na^+^/H^+^ or K^+^/H^+^ exchange at minimal salt concentration, 0.5 mM of (NaCl or KCl). NhaC1 and NhaC2 were moderately active at pH 9.5 but less efficient than NhaC3 (Fig 1). In table 1, *K*_*m*_ values for catalytic activities of NhaC1, NhaC2 and NhaC3 were calculated at optimum pH 9.5 using *E. coli* as a host.

**Figure 1:**
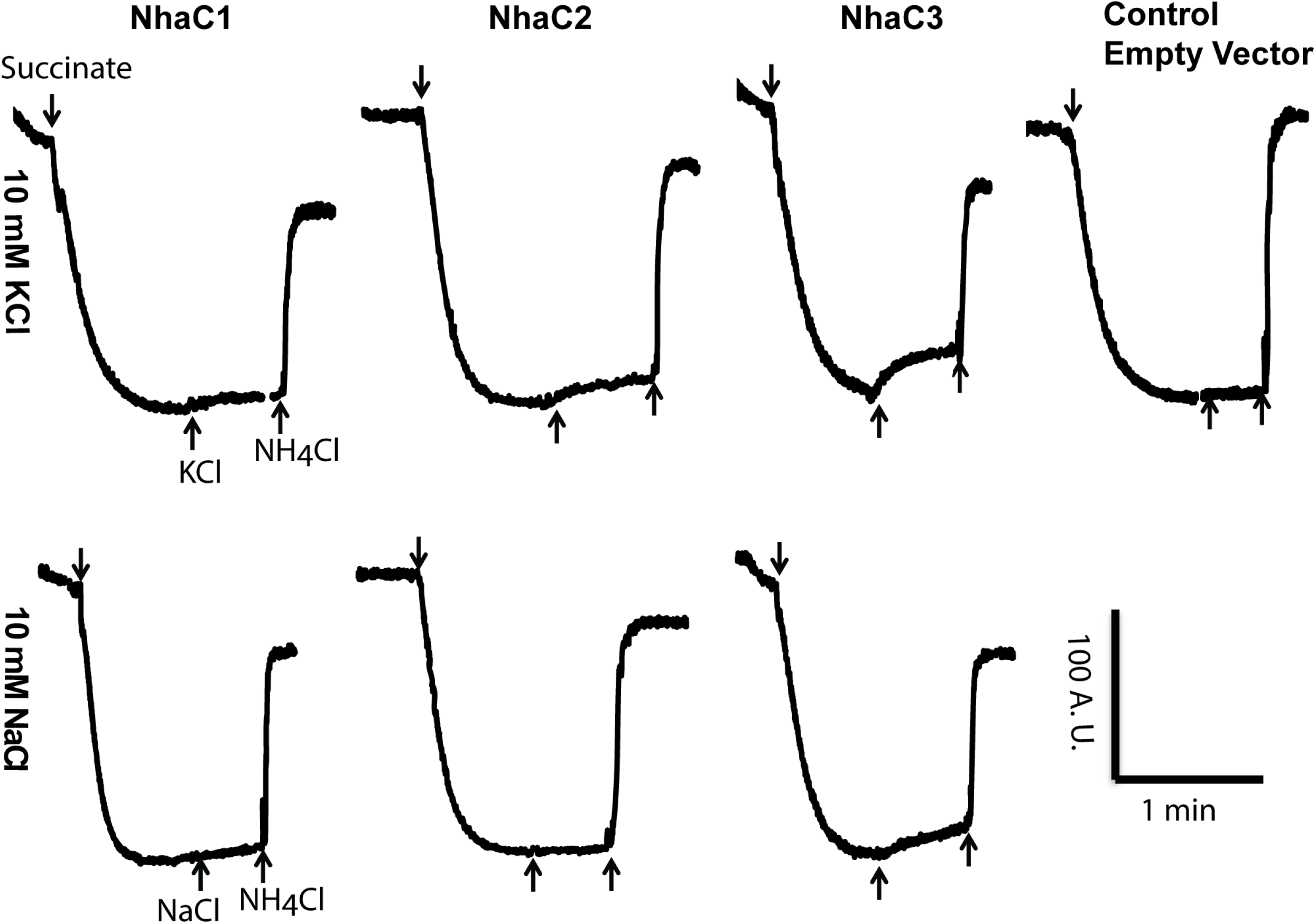
Antiporter activity of NhaC1, NhaC2 and NhaC3 were assayed at pH 9.5 for Na^+^/H^+^ and K^+^/H^+^ exchange. Empty pBAD vector in first lane was taken as control for measuring Na^+^/H^+^ and K^+^/H^+^ exchange. The everted membrane vesicles were prepared using *E. coli* KNabc host transformed with inducible pBAD vector in which *S. aureus nhaC* genes were cloned. Addition of 2.5 mM succinate in assay buffer generated PMF which was monitored as florescence quenching using 1 M acridine orange as ΔpH probe. The antiport activity was measured as percentage dequenching in florescence after adding 10 mM of NaCl or KCl. 1 mM NH_4_Cl was used to terminate the reaction which establish a baseline. The tracing is representative of antiport assay carried out with three independent vesicle preparations and conducted in duplicate for each preparation. A. U. is arbitrary units.

**Table 1:** NhaC1, NhaC2 and NhaC3 antiporter activities at pH 9.5^a^.

| Antiporter | Substrate<br>(10 mM) | Maximum activity at pH 9.5<br>(% Dequench) | $K_m$ value<sup>b</sup><br>(mM) |
| --- | --- | --- | --- |
| NhaC1 | K<sup>+</sup> | 13 | 0.42±0.16 |
|  | Na<sup>+</sup> | 10 | - |
| NhaC2 | K<sup>+</sup> | 10 | 0.75±0.04 |
|  | Na<sup>+</sup> | 5 | - |
| NhaC3 | K<sup>+</sup> | 36 | 0.45±0.03 |
|  | Na<sup>+</sup> | 30 | 0.42±0.02 |

<sup>a</sup>Antiporter assay was conducted using 2.5 mM tris-succinate to generate proton motive force feeding proton inside the everted vesicles. The everted membrane vesicles were prepared using *E. coli* KNa bc host transformed with inducible pBad vector in which *S. aureus nhaC1*, *nhaC2* and *nhaC3* were cloned along with empty vector as control. Assays were conducted between pH values 7.0 to 9.5 (data not shown). NhaC candidates were active at pH 9.5 exhibiting maximum dequenching.
<sup>b</sup> $K_m$ values ( ± standard deviation) were calculated as average of three independent set of experiment that was conducted in duplicate with freshly prepared membrane vesicles from pregrown culture.

Similar to NhaCs, CPA1 candidates actively exchanged Na^+^/H^+^ and K^+^/H^+^ between pH values 7 to 9.5. In Fig 2, raw data of antiport activities indicates that CPA1-1 exhibited antiporter activity near neutral pH whereas CPA1-2 was active at larger range of pH between 7 −9.5 and had robust activity > 93% dequenching at pH 9.0 in presence of minimal salt concentration of 0.5 mM of Na^+^ or K^+^ salt, exhibiting highest activity among whole cohort of antiporters. The antiport activity profile is mentioned in Table 2 which includes *K*_*m*_ values of each CPA1-1 and CPA1-2 at their optimal pH, 7.5 and 9.0 respectively. CPA2 did not exhibit any antiport activity for Na^+^ or K^+^/H^+^ exchange within the pH range 7-9.5 (data not shown). This indicates the possibility that CPA2 might have catalytic activity other than antiporter.

**Figure 2:**
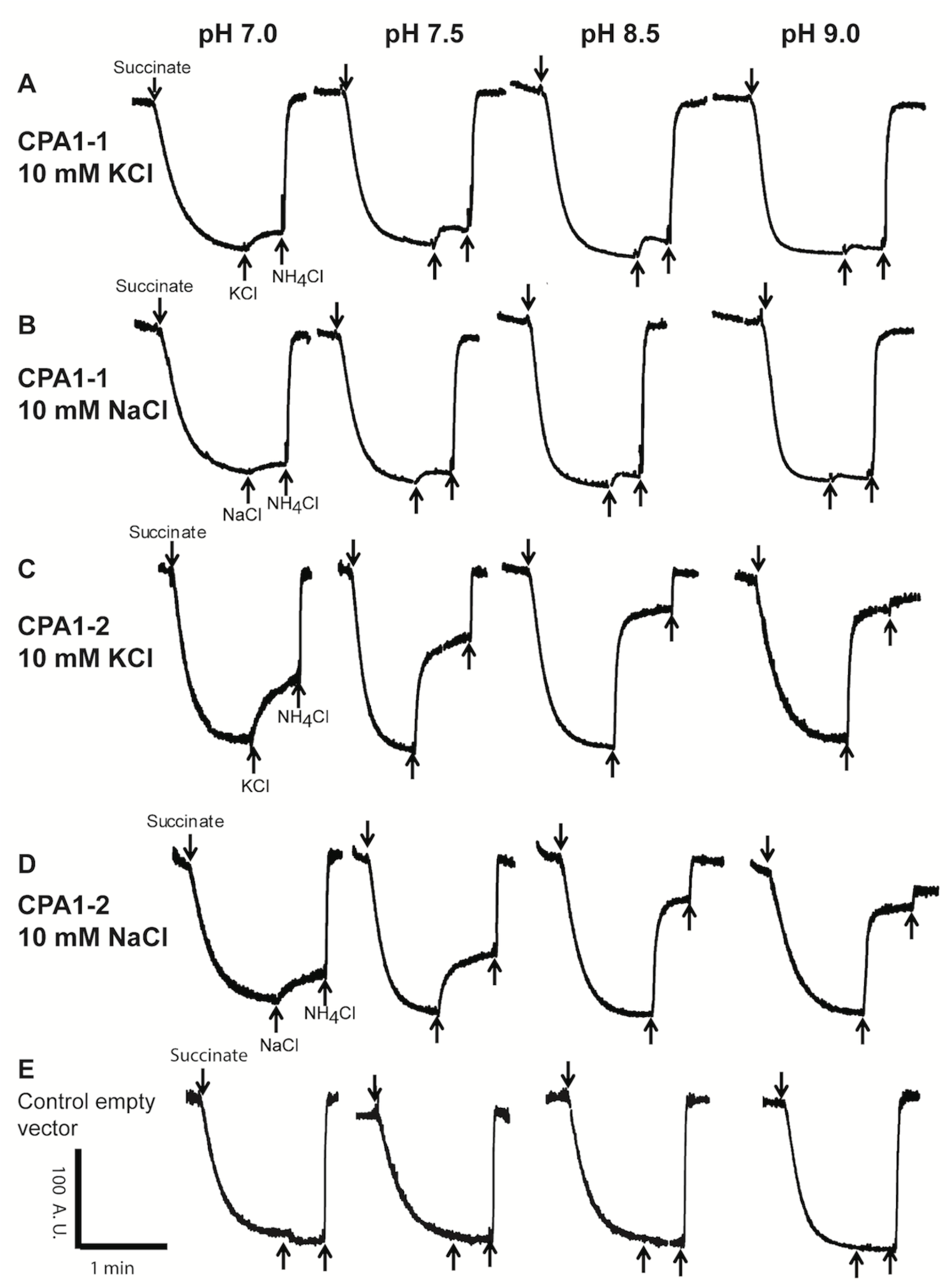
Antiporter activity of CPA1-1(A and B) and CPA1-2 (C and D) were performed as function of pH for Na^+^/H^+^ and K^+^/H^+^ exchange. Lane (E) is representative of control assay performed using Na^+^ and K^+^ salt resulting in no dequenching with change in pH. *S. aureus* Newman *cpa1-1 and cpa1-2* genes were overexpressed into inducible pBAD vector and transformed into *E. coli* KNabc host. These strains were used for large scale culture and preparing membrane vesicles for the assay. Addition of 2.5 mM succinate in assay buffer generated PMF which was monitored as florescence quenching using 1 M acridine orange as ΔpH probe. The antiport activity was measured as percentage dequenching in florescence after adding 10 mM of NaCl or KCl. 1 mM NH_4_Cl was used to terminate the reaction, which establish a baseline. The tracing is representative of antiport assay carried out with three independent vesicle preparations and conducted in duplicate for each preparation. A. U. is arbitrary units.

**Table 2:** CPA1-1 and cPA1-2 activity profile at optimal pH^a^.

| Antiporter | Substrate<br>(10 mM) | Optimal assay<sup>b</sup><br>pH | Maximum activity at<br>assay pH (%dequench) | $K_m$ Values<sup>c</sup><br>(mM) |
| --- | --- | --- | --- | --- |
| CPA1-1 | K<sup>+</sup> | 7.5 | 16 | 14.81±11.16 |
|  | Na<sup>+</sup> | 7.5 | 10 | 3.59±0.36 |
| CPA1-2 | K<sup>+</sup> | 9.0 | 75 | 3.85±0.72 |
|  | Na<sup>+</sup> | 9.0 | 82 | 2.96±0.32 |

<sup>a</sup>Antiporter assays were performed in everted membrane vesicles using *E. coli* cells which were transformed with inducible pBad vector with cloned *S. aureus cpa1-1* and *cpa1-2* genes and taking empty vector controls. 2.5 mM tris-succinate was used in assay buffer to energize respiratory chain generating PMF.
<sup>b</sup>Assay were conducted on pH range 7.0 to 9.5 using NaCl and KCl. Assay pH on which CPA1-1 and CPA1-2 exhibited maximum dequenching was considered as optimal.
<sup>c</sup> $K_m$ values ( ± standard deviation) were calculated as average of three independent set of experiment that was conducted in duplicate with freshly prepared membrane vesicles from pregrown culture.

### Comparison of physiological roles of NhaC, CPA1 and CPA2 antiporters under stress conditions

In order to explore the contribution of NhaC candidates in *S. aureus* growth and physiology, single, double and triple knockout strains: *ΔnhaC1, ΔnhaC2* and *ΔnhaC3*; *ΔnhaC1ΔnhaC3* and *ΔnhaC1ΔnhaC2ΔnhaC3* were constructed in Newman. Growth experiments were conducted in Luria-Bertani broth medium with and without supplemented salt conditions (LB0). None of the mutants showed growth defect at pH 7.5 and 8.5 in absence of added salt and, at pH 9.5 *ΔnhaC3*, *ΔnhaC1ΔnhaC3* and *ΔnhaC1ΔnhaC2ΔnhaC3* exhibited significant growth deficit (Fig 3A). When growth media was supplemented with 1 M of Na^+^ or K^+^ salt, *ΔnhaC3*, double and triple knockouts were found to be sensitive at pH 8.5. The complete loss of viability was observed in all knockouts at pH 9.5 supplemented with 1 M of sodium salt (Fig 3A). This result was consistent with catalytic properties of NhaC antiporters at pH 9.5 and indicated their major role in alkali tolerance of *S. aureus.*

**Figure 3:**
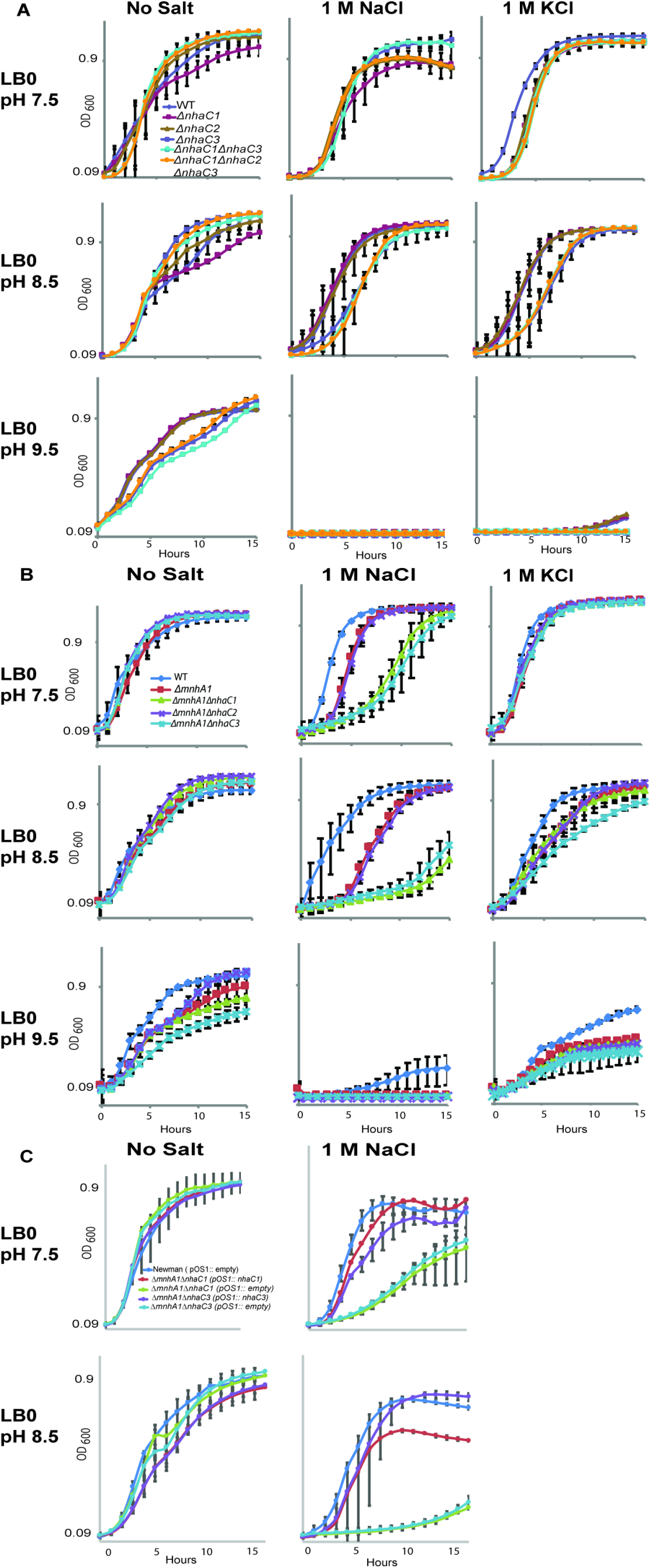
(A) Physiological role of NhaC1, NhaC2 and NhaC3 were assessed on single, double and triple deletions of *nhaC* genes in Newman at pH 7.5, 8.5 and 9.5 using 1 M NaCl and 1 M KCl. In lane B, double knockouts strains of *mnh1* with *nhaC1* and *nhaC3* respectively were used under same the condition. For growth experiments shown in all panels, strains were grown in pH 7.5 LB0 at 37°C. 1 M NaCl and 1 M KCl were added as indicated. Lane C represents complementation of wild type Newman *nhaC1* and *nhaC3* genes into *ΔnhaC1ΔmnhA1* and *ΔnhaC3ΔmnhA1* knockout strains using pOS1 vector and using pOS1::empty in respective double knockouts as a control. The growth curve assay was conducted in three independent sets of experiments. Error bars represent the standard deviation.

In the previous study (18) it was reported that Mnh1 significantly contributes in pH homeostasis at pH 7.5 and Mnh2 was active at higher pH range. However, it was still unclear that how rest of the antiporters in cohort regulates pH tolerance and salt tolerance in extreme conditions. Therefore, it was interested to investigate how *S. aureus* regulates pH homeostasis by NhaCs that exhibit catalytic activity at alkaline pH and orchestrate with Mnh1 that exchange Na^+^/H^+^ at lower pH under stress conditions. In order to explore this phenomenon, *nhaC1, nhaC2* and *nhaC3* were deleted with *mnhA1* respectively to construct double mutant strains. Severe growth defects were observed in *ΔnhaC1ΔmnhA1*, *ΔnhaC2ΔmnhA1*, and *ΔnhaC3ΔmnhA1* knockouts in various stress conditions. Increase in pH exacerbated growth defects with supplemented sodium or potassium salt. By the addition of 1 M sodium salt in growth medium at pH 7.5, growth defects were significantly higher in *ΔnhaC1ΔmnhA1* and *ΔnhaC3ΔmnhA1* in comparison to *ΔmnhA1* or *ΔnhaC1 and ΔnhaC3* alone (Fig 3B). Increasing pH to 8.5 increased the severity of the growth deficit in these double knockouts. Complementation of *nhaC1* and *nhaC3* genes into double mutant strains using pOS1 vector, restored the growth of these mutants at *ΔmnhA1* level under tested conditions (Fig 3C). The data shown above clearly indicate that Mnh1 and NhaCs synergistically play an important role to support *S. aureus* in coping sodium toxicity and alkaline pH.

In addition to explore the role of CPA1 family candidates in *S. aureus* physiology, single knockout of *cpa1-1* and *cpa1-2*; *Δcpa1-1 and Δcpa1-2* and double knockout; *Δcpa1-1Δcpa1-2* were constructed in Newman. Single knockouts of *cpa1-1* and *cpa1-2* have mild growth defects at pH 7.5 and 8.5; however, double deletion of both genes together had severe consequence in pH tolerance of *S. aureus* (Fig 4A). None of the mutant strains, single or double knockouts of CPA1 were found sensitive to sodium or potassium stress at pH 7.5 and 8.5. This data indicates that salt tolerance of *S. aureus* was not affected at pH 7.5-8.5, in absence of CPA1 family candidates, specially CPA1-2, which exhibited highest antiport activity at wider pH range, and the cation/proton exchange was efficiently overtaken by another antiporter in cohort (Fig 7). To find the synergistic relationship between CPA1 candidates and Mnh1, *cpa1-1* and *cpa1-2* were deleted with *mnhA1* respectively to construct double knockouts. In absence of supplemented Na^+^ or K^+^ salt, *Δcpa1-1ΔmnhA1* and *Δ*cpa1-*2ΔmnhA1* exhibited severe growth defect at pH 7.5 and the severity of growth defect was further enhanced at pH 8.5. Addition of 1 M Na^+^ salt in growth medium at pH 7.5 exhibited severe consequence on growth deficit in *Δcpa1-1ΔmnhA1* and *Δ*cpa1-*2ΔmnhA1* which got worsened at pH 8.5 (Fig 4B). The growth deficit in double knockouts were reversed upto *ΔmnhA1* level by complementing *cpa1-1* and *cpa1-2* genes in their respective double knockout strains (Fig 4C). There was no growth deficit reported in *Δcpa2* mutant strain at any stress conditions indicating no obvious role of CPA2 antiporter in pH or salt tolerance (Fig 4A).

**Figure 4:**
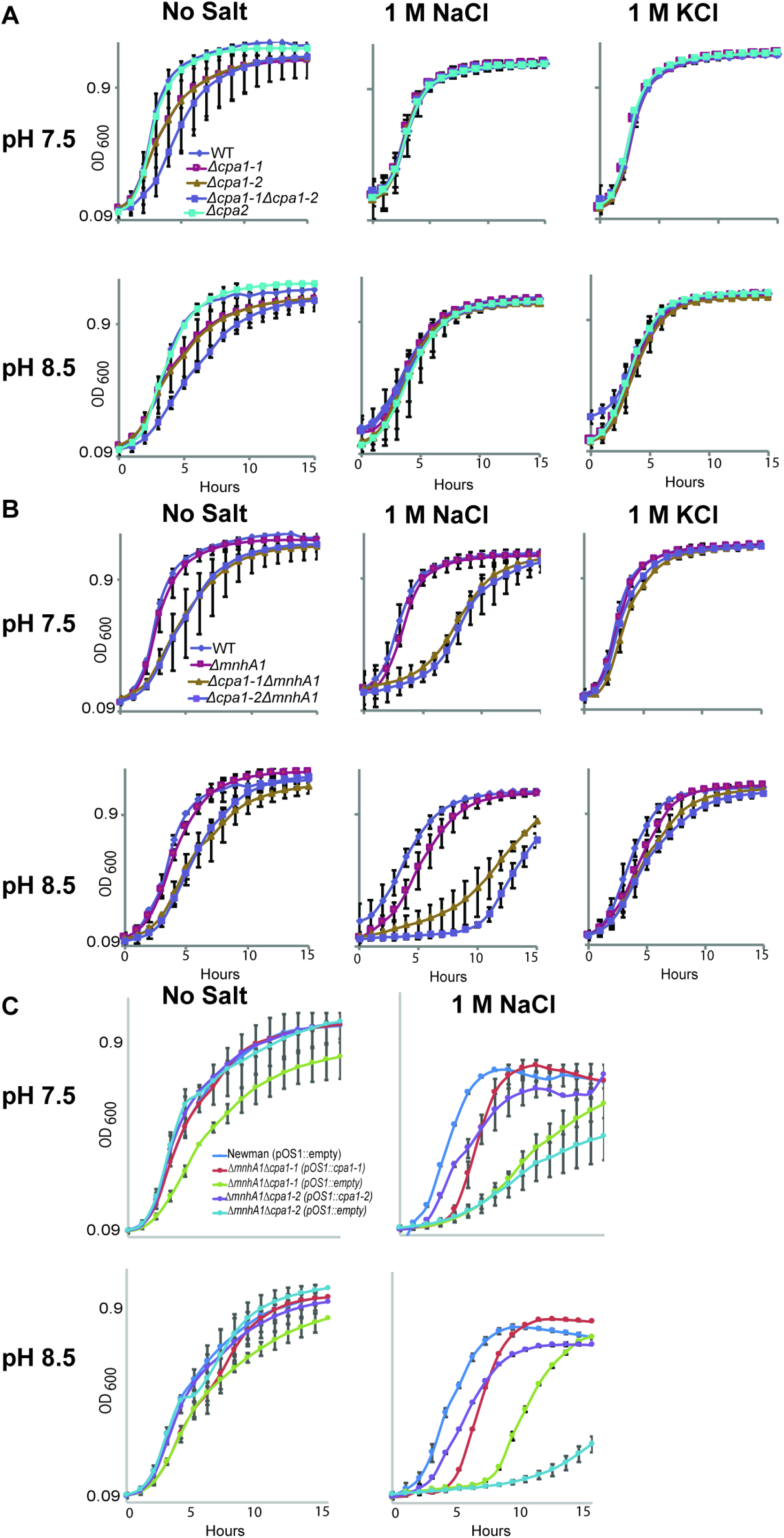
(A) Physiological role of CPA1-1, CPA1-2 and CPA2 were assessed with single, and double deletions of *cpa1-1, cpa1-2 and cpa2* genes in Newman at pH 7.5, 8.5 and 9.5 using 1 M NaCl and 1 M KCl. In lane B, double knockouts strains of *mnh1* with *cpa1-1* and *cpa1-2* respectively, were used under same the condition. For growth experiments shown in all panels, strains were grown in pH 7.5 LB0 at 37°C. 1 M NaCl and 1 M KCl were added as indicated. (C) For complementation experiments, wild type Newman *cpa1-1* and *cpa1-2* were complemented in respective double deletion strains of *Δcpa1-1ΔmnhA1* and *Δcpa1-2ΔmnhA1* using pOS1 vector. Double knockouts transformed with pOS1::empty were taken as control strains. The growth curve assay was conducted in three independent sets of experiments. Error bars represent the standard deviation.

### Expression profiles of NhaC, CPA1 and CPA2 antiporters as function of pH

We compared expression level of three NhaCs, two CPA1, two CPA3 (Mnh1 and Mnh2) and CPA2 antiporters grown in LB0 media without any supplemented Na^+^ or K^+^ salt using culture at log phase. The wild type Newman strain was grown in pH 6.0, 7.5, and 9.5 to compare the fold change of antiporter genes in acidic, neutral and alkaline pH conditions. CPA1-2, Mnh2, NhaC1, NhaC2, and NhaC3 exhibited higher expression at pH 9.5 and expression of NhaC3 was highest among all eight CPAs. The expression profile of these antiporters was consistent with their antiporter activities at alkaline pH. In contrast, Mnh1 and CPA1-1 were expressed at acidic side of pH. CPA2 did not show any change in expression level at acidic or alkaline pH (6.0 and 9.5) (Fig 5). These data sets were analyzed using quantitative PCR assay on each of eight antiporters is *S. aureus* Newman.

**Figure 5:**
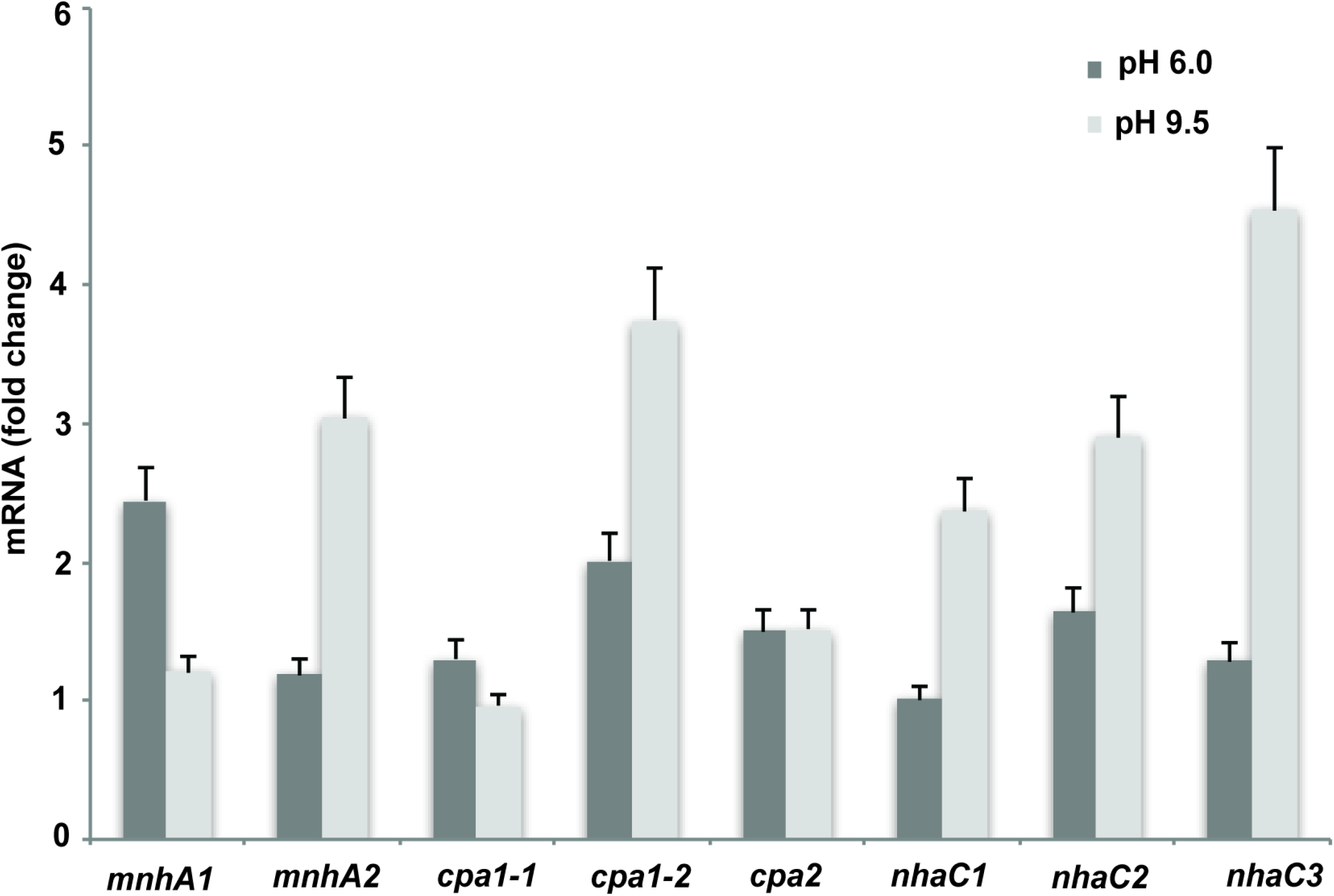
Expression of *cpa1-1, cpa1-2, cpa2, mnhA1, mnhA2, nhaC1, nhaC2*, and *nhaC3* genes in *S. aureus* Newman were assessed at pH 6.0 and pH 9.5 without any added salt. *Pyk, fabD* and *QoxB* were used as reference genes. Data represent average of biological triplicates and error bars show represent standard deviation.

### Importance of NhaC and CPA1 antiporters in pathogenesis of *S. aureus*

The contribution of NhaC1, NhaC2, NhaC3, CPA1-1 and CPA1-2 in *S. aureus* pathogenesis was evaluated by systemic infection of isogenic mutant strain using murine model and Newman wild type as a control of virulence. Strains with *Δcpa1-1, Δcpa1-2, ΔnhaC1* and *ΔnhaC2* deletions exhibited virulence similar to Newman wild type, whereas strain with *ΔnhaC3* deletion showed marked attenuation of virulence (Fig 6A). To confirm the virulence defect observed in *ΔnhaC3* strain is due to lack of functional *nhaC3*, complemented strain of wild type *nhaC3* in *ΔnhaC3* mutant strain was constructed by using pOS1 plasmid as mentioned in previous studies (18, 19). The virulence defect was reversed with the restoration of wild type *nhaC3* in *ΔnhaC3* mutant strain (Fig 6B). The survival data shown in Fig 6C was consistent with ~ 3.3-log reduction in bacterial burden observed in kidneys of mice infected with *ΔnhaC3* as compared to mice infected with wild type *nhaC3*.

**Figure 6:**
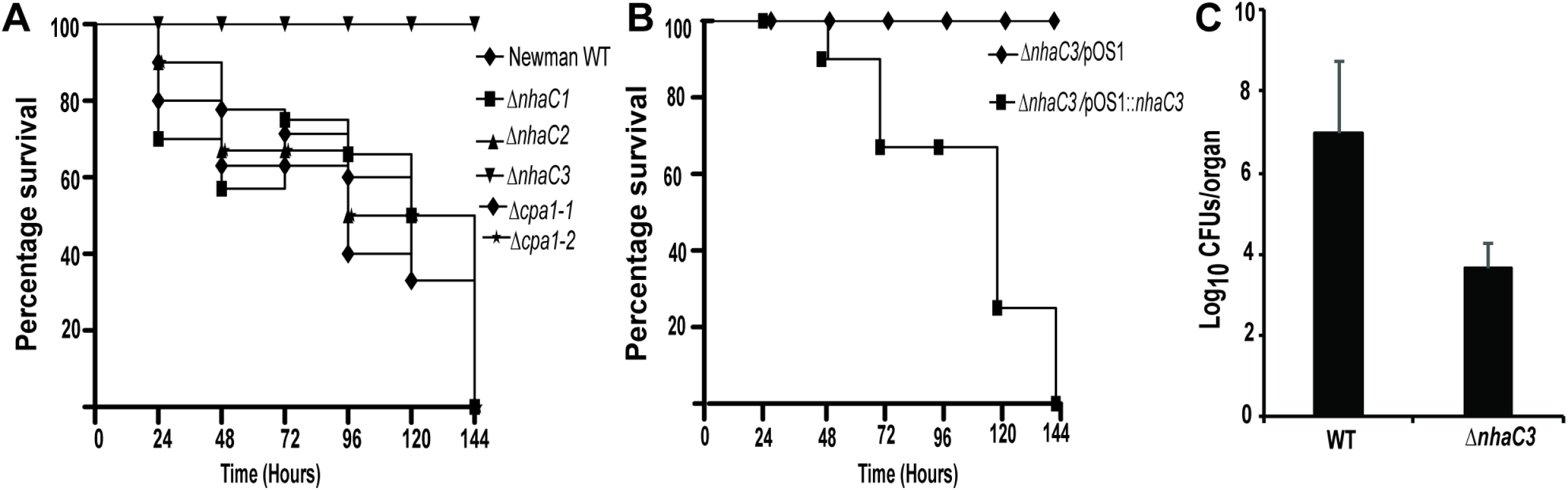
*In-vivo* assay for *S. aureus* pathogenesis. **(A)** Survival curve of mice n=10 in each group was infected intravenously using ~1×10^7^ CFU of Newman wild type, Δ*cpa1-1*, Δ*cpa1-2*, Δ*nhaC1*, Δ*nhaC2* and Δ*nhaC3.* **(B)** Survival curve of mice infected intravenously with Δ*nhaC3*/pOS1::empty (n=10) and Δ*nhaC3*/pOS1::*nhaC3* (n=10). **(C)** CFU count from kidney tissues harvested from mice infected with Newman wild type and Δ*nhaC3* after ~96 hours of post infection. The error bar represents standard deviation.

## Methods and Materials

### Bacterial strains, plasmids and primers

The bacterial strains, plasmids and primers used in this study are listed in Table 3 and 4.

**Table 3:**
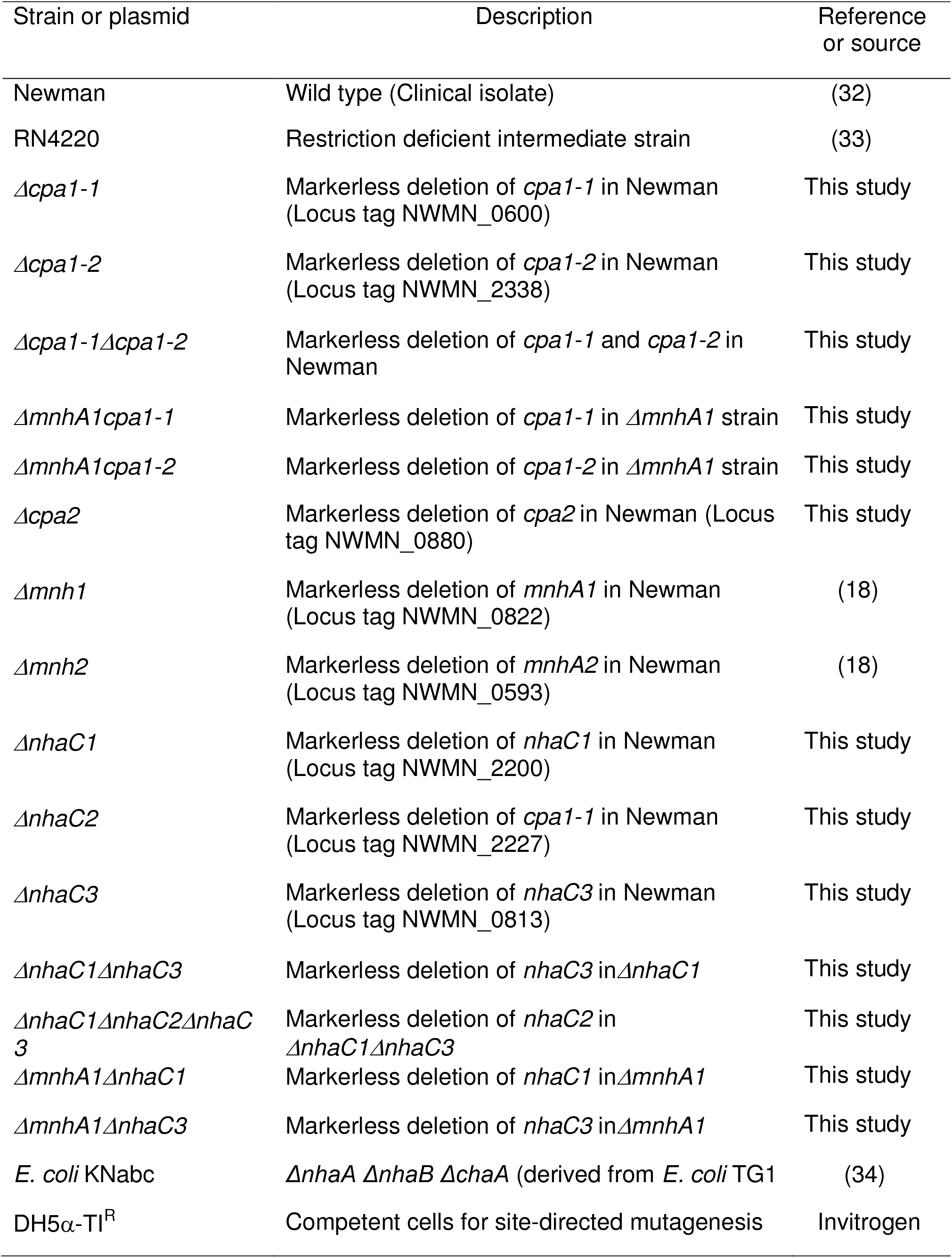
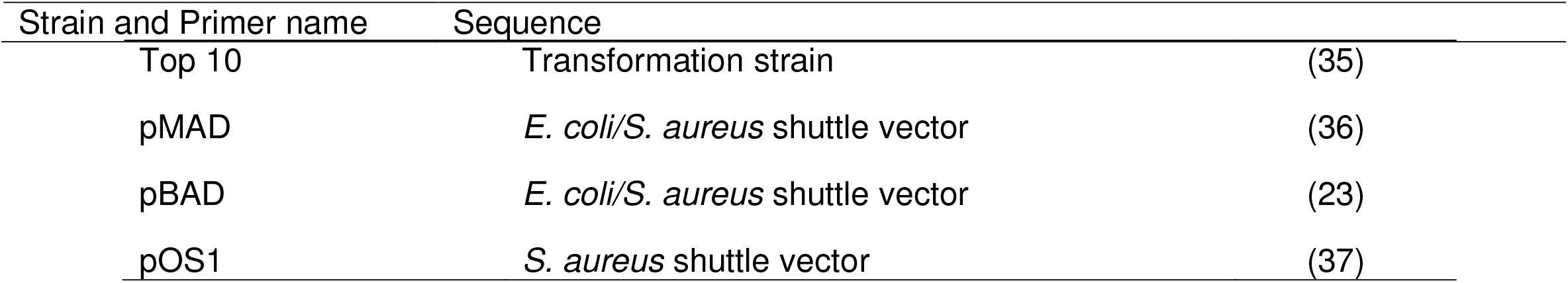
Bacterial strains, plasmids and primers used in this study.

| Strain or plasmid | Description | Reference or source |
| --- | --- | --- |
| Newman | Wild type (Clinical isolate) | (32) |
| RN4220 | Restriction deficient intermediate strain | (33) |
| $\Delta cpa1-1$ | Markerless deletion of *cpa1-1* in Newman (Locus tag NWMN_0600) | This study |
| $\Delta cpa1-2$ | Markerless deletion of *cpa1-2* in Newman (Locus tag NWMN_2338) | This study |
| $\Delta cpa1-1\Delta cpa1-2$ | Markerless deletion of *cpa1-1* and *cpa1-2* in Newman | This study |
| $\Delta mnhA1cpa1-1$ | Markerless deletion of *cpa1-1* in $\Delta mnhA1$ strain | This study |
| $\Delta mnhA1cpa1-2$ | Markerless deletion of *cpa1-2* in $\Delta mnhA1$ strain | This study |
| $\Delta cpa2$ | Markerless deletion of *cpa2* in Newman (Locus tag NWMN_0880) | This study |
| $\Delta mnh1$ | Markerless deletion of *mnhA1* in Newman (Locus tag NWMN_0822) | (18) |
| $\Delta mnh2$ | Markerless deletion of *mnhA2* in Newman (Locus tag NWMN_0593) | (18) |
| $\Delta nhaC1$ | Markerless deletion of *nhaC1* in Newman (Locus tag NWMN_2200) | This study |
| $\Delta nhaC2$ | Markerless deletion of *cpa1-1* in Newman (Locus tag NWMN_2227) | This study |
| $\Delta nhaC3$ | Markerless deletion of *nhaC3* in Newman (Locus tag NWMN_0813) | This study |
| $\Delta nhaC1\Delta nhaC3$ | Markerless deletion of *nhaC3* in $\Delta nhaC1$ | This study |
| $\Delta nhaC1\Delta nhaC2\Delta nhaC3$ | Markerless deletion of *nhaC2* in $\Delta nhaC1\Delta nhaC3$ | This study |
| $\Delta mnhA1\Delta nhaC1$ | Markerless deletion of *nhaC1* in $\Delta mnhA1$ | This study |
| $\Delta mnhA1\Delta nhaC3$ | Markerless deletion of *nhaC3* in $\Delta mnhA1$ | This study |
| *E. coli* KNabc | $\Delta nhaA \Delta nhaB \Delta chaA$ (derived from *E. coli* TG1) | (34) |
| DH5α-TI<sup>R</sup> | Competent cells for site-directed mutagenesis | Invitrogen |

| Strain and Primer name | Sequence |  |
| --- | --- | --- |
| Top 10 | Transformation strain | (35) |
| pMAD | *E. coli/S. aureus* shuttle vector | (36) |
| pBAD | *E. coli/S. aureus* shuttle vector | (23) |
| pOS1 | *S. aureus* shuttle vector | (37) |

**Table 4:**
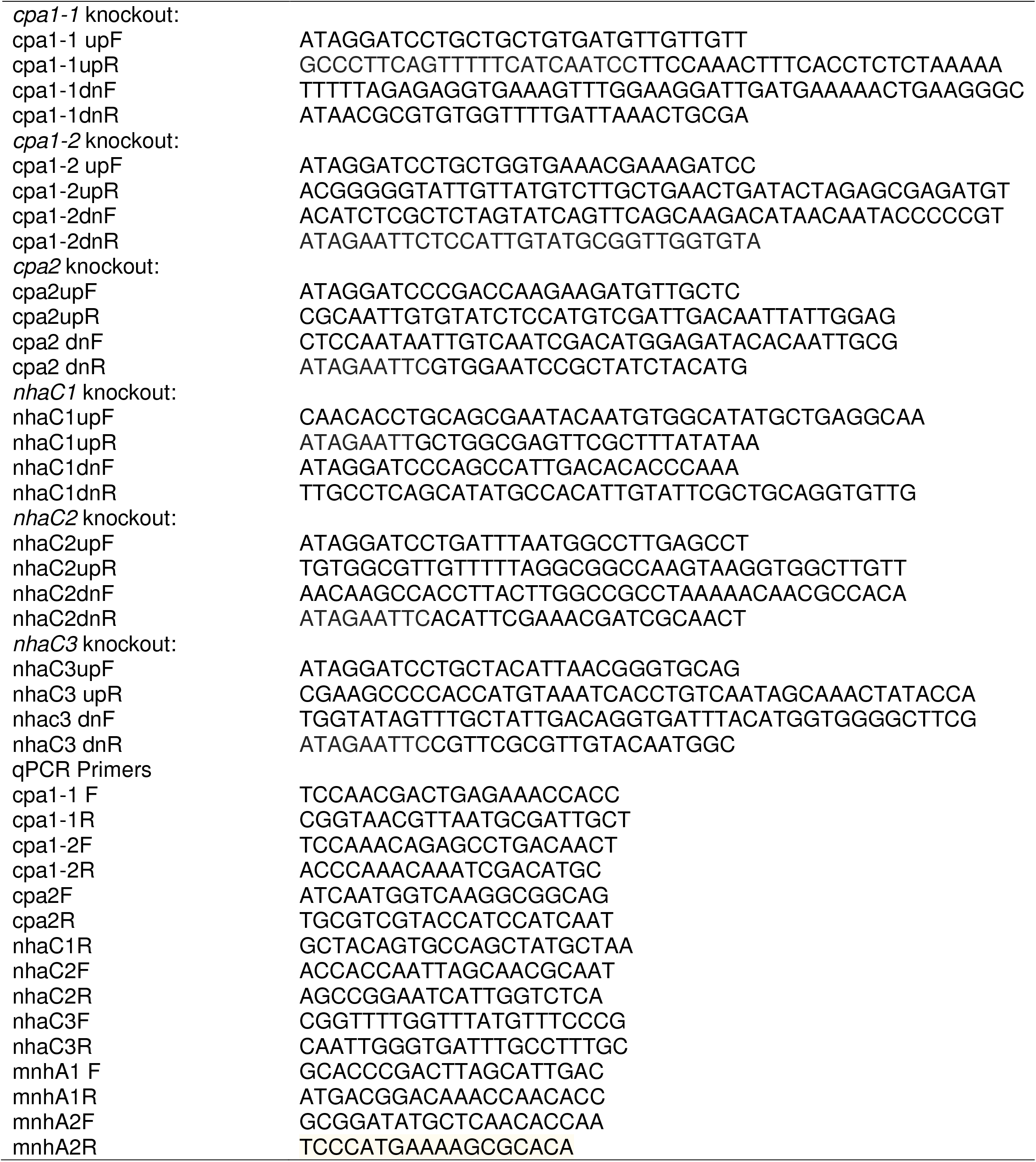
Primers used in this study.

### Growth conditions

*S. aureus* strains were routinely grown in a modified version of Luria-Bertani broth (LB), designated as LB0, which is LB without added NaCl or KCl. Cultures from frozen stock were incubated at 37°C with shaking at 225 rpm. Erythromycin at 2.5 μg/ml, and chloramphanicol 10 μg/ml were added in the medium to grow *S. aureus* for various plasmid selections.

### Growth curve experiments

Growth experiments were done according to our previous study (18). Briefly, glycerol stocks of *S. aureus* wild type and mutant strain were inoculated in LB0 medium, pH 7.5, and grown for 16 h at 37°C with shaking at 225 rpm prior to growth experiments. Cultures were grown overnight and normalized to an OD_600_ of 0.2 with unbuffered LB0 medium. Ten micro liters of pre-cultures at an OD_600_ of 0.2 was passed into 190 μl of corresponding medium in 96-well microplates starting with OD_600_ of 0.01 for all growth conditions. Microplate lids were then carefully sealed with 1.2 × 40 cm silicone rubber tape and incubated at 37°C with shaking at 225 rpm in a BioTek Power Wave HT microplate spectrophotometer for 24 h. OD_600_ readings were collected every hour. Growth curves were calculated as averages of at least three independent experiments done in triplicate repeats.

### Construction of marker less deletions in *S. aureus* by allelic replacement

In frame deletions of target genes were generated by using pMAD according to previously published methods (20). Briefly, ∼1-kb PCR products on either side of the sequence to be deleted were generated and fused by gene SOEing (21). 2-kb product was ligated into pMAD and transformed into *E. coli*. After plasmid isolation and sequence verification, the construct was moved into *S. aureus* RN4220 by electroporation. After isolation from RN4220, the construct was electroporated into the target *S. aureus* strain. The plasmid was recombined into the chromosome by inoculating a liquid culture for 2 h at the permissive temperature (28°C), followed by overnight inoculation at the restrictive temperature (42°C) and plating of dilutions on LB0 agar containing erythromycin. Merodiploid clones (containing the plasmid recombined into the chromosome) were verified by PCR. To resolve the plasmid out of the chromosome and to generate candidate deletion mutants, liquid cultures of merodiploids were incubated at 28°C without selection and transferred by 1:100 dilutions for 7 days before being plated on LB0 agar. Candidate mutants were screened for loss of erythromycin resistance (confirming loss of the plasmid), and PCR and sequencing was used to confirm exclusive presence of the deleted allele.

### Preparation of the total RNA and cDNA from *S. aureus* Newman and relative quantification of RNA transcripts by qPCR

RNA was prepared according to a method described previously (22). Newman wild type was inoculated in 50 ml of LB0 medium and grown up to an OD_600_ of 0.9 at 37°C. Briefly, eight milliliters of culture was added to 8 ml of RNA Protect bacteria reagent (Qiagen) in 50-ml sterile tubes and then vortexed immediately for 5 sec and incubated at room temperature for 5 min. Cells were harvested by centrifugation (4,700 rpm, 21°C, and 10 min), the supernatant was poured off, and then the tube was inverted on paper towel for 10 sec. Pellets were stored at −80°C overnight. The following day, RNA was isolated using an miRNeasy purification kit (Qiagen) for subsequent steps as described in (Vaish et al 2018). Samples were run in triplicate, and a no-template control and a no-reverse transcriptase control were run to ensure absence of DNA contamination. Primers used in this study are mentioned in table 4. Data were analyzed using SDS, version 2.2.1, software (Applied Biosystems, USA).

### Antiporter assays

Antiporter assays were conducted in everted membrane vesicles prepared from transformants of the triple antiporter-deficient *E. coli* KNabc strain expressing the empty vector, pBAD, or Newman *cpa1-1, cpa1-2, cpa2, nhaC1, nhaC2 and nhaC3* genes. The gene of interest was placed into the pBAD vector downstream of the araBAD promoter, which drives expression of the gene of interest in response to 0.002% L-arabinose added in early exponential phase of the culture (23). The everted membrane vesicles are oriented in such a manner that part of the membrane that is exposed outside the bacterial cells comes inside the vesicles. The transformants were grown for 3 hours after inducing with 0.002% arabinose and then frozen in liquid nitrogen and stored at −80°C. Preparation of vesicles from the *E. coli* transformants was conducted using a French press as described earlier (24). The vesicles were used immediately after preparation, without being frozen. The assays also followed a protocol used earlier with the same buffer and pH conditions. Acridine orange was used as the ΔpH probe. The measurement was conducted using an RF-5301 PC Shimadzu Spectro fluorophotometer equipped with a stirrer, with excitation at 420 nm and emission at 500 nm (both with a 10-mm slit). When the respiratory chain is energized by succinate, the respiratory chain starts pumping protons inside the vesicles (as vesicles are everted). The initiation of proton motive force generation is indicated for specific experiments.

### Complementation of markerless deletions

To complement *ΔmnhA1Δcpa1-1*, *ΔmnhA1Δcpa1-2*, *ΔmnhA1ΔnhaC1*, *ΔmnhA1ΔnhaC3* and *ΔnhaC3* with wild type Newman *cpa1-1, cpa1-2, nhaC1* and *nhaC3* respectively, genes were amplified by PCR and inserted into pOS1 vector (25). Briefly, cpa*1-1, cpa1-2, nhaC1* and *nhaC3* were amplified using primers mention in table 4 and then ligated into pOS1 vector at their respective restriction sites resulting in transformation in *E. coli* using ampicillin for selecting colonies. Plasmid construct were isolated and sequenced verified. After that, these constructs were transformed into *S. aureus* RN4220 by electroporation followed by electroporating constructs in appropriate target mutants of *S. aureus* Newman.

### Murine systemic infection

Animal experiments were performed by protocol approved by Mount Sinai Institutional Animal Care and Use Committee. For systemic infections, 5 weeks old female swiss webstar mice, N=10, were injected with 1×10^7^ CFU/ml intravenously in tail using Newman wild type, isogenic mutant strains of *Δcpa1-1*, *Δcpa1-2*, *ΔnhaC1*, *ΔnhaC2* and *ΔnhaC3* and complemented strain *ΔnhaC3/pOS1::nhaC3* (Fig 6A,B). Mice were monitored for acute infection and sign of morbidity upto seven days and survival curve were plotted over time using Prism software. Bacterial load experiment was done on kidney tissues harvested ~96 hours of post infection and euthanizing mice with CO_2_ when acute infection signs appears (Fig 6C).

## Discussion

*S. aureus* is a neutralophile and can act like an alkaliphile in extreme conditions inside host as well as in the environment. The viability of *S. aureus* at alkaline pH highly depends on secondary transporters, which maintain inside cytoplasmic pH near 7.5-7.7. This study confirms that *S. aureus* NhaC and CPA1 family candidates are secondary antiporters that are involved in Na^+^ and/or K^+^ efflux in exchange of protons. The existences of NhaC type antiporters are prevalent in large groups of pathogenic and non-pathogenic bacterial species as such as *L. hongkongensis, C. violaceum, N. meningitidis, N. gonorrhoeae, Heamophilus influenzae* shown to have important role in alkali-tolerance (26-28). Ivey DM et. al. (3) observed the role of NhaC for the first time in non-pathogenic *B. pseudofirmus*. We investigated the physiological role of NhaCs in highly resistant pathogen *S. aureus* that successfully survives in alkaliphilic conditions in the environment as well as human host. All three *S. aureus* NhaCs are structurally distinct and shows structural identity <27% on alignment tool. Interestingly, NhaC3 showed robust activity among three NhaCs and also exhibited significantly distinct synergistic relationship with Mnh1 in halotolerance. Since Mnh1 regulates sodium toxicity and pH homeostasis at pH 7.5 and NhaCs are active at high alkaline pH 9.5, it was interesting to observe the effect of both types of antiporters on overall pH regulation of *S. aureus* under wide range of osmotic and pH stresses. We constructed double knockouts of *nhaC1, nhaC2*, and *nhaC3* with *mnh1* respectively. When these double knockouts were grown at pH range 7.5 to 9.5, severe consequence of alkaline stress was observed at pH 9.5. Supplementing growth medium with 1 M NaCl made *ΔnhaC1Δmnh1* and *ΔnhaC3Δmnh1* double knockouts highly sensitive even at pH 7.5 that further exacerbated with increasing pH. Altogether, these findings confirm the cross talk between NhaC1, NhaC3, and Mnh1 in regulating pH and maintaining cell homeostasis.

Similarly, the results confirm that both of CPA1 candidates actively exchange Na^+^/H^+^ and K^+^/H^+^ between pH ranges 7 to 9.5 (Fig 2). CPA1-1 catalyzes Na^+^/K^+^ and H^+^ exchange near neutral pH 7-8. While CPA1-2 is active within a wide range of pH and actively maintains homeostasis at alkaline conditions. CPA1-2 exhibited maximum dequenching which reaches upto ~93% among whole cohort of CPA in *S. aureus* indicating its active involvement in pH homeostasis in alkaline environment. However, *Δcpa1-2* did not show any significant decrease in growth pattern under sodium toxicity (Fig 4) possibly due to compensatory activation of other antiporters at low (Mnh1) as well as high pH (NhaCs and Mnh2) (Fig 7). In *Δcpa1-2Δmnh1*, growth deficit was significantly higher under stress conditions than single knockouts of both genes, which clearly indicates the synergistic functional overlapping of *cpa1-2* with *mnh1*.

**Figure 7:**
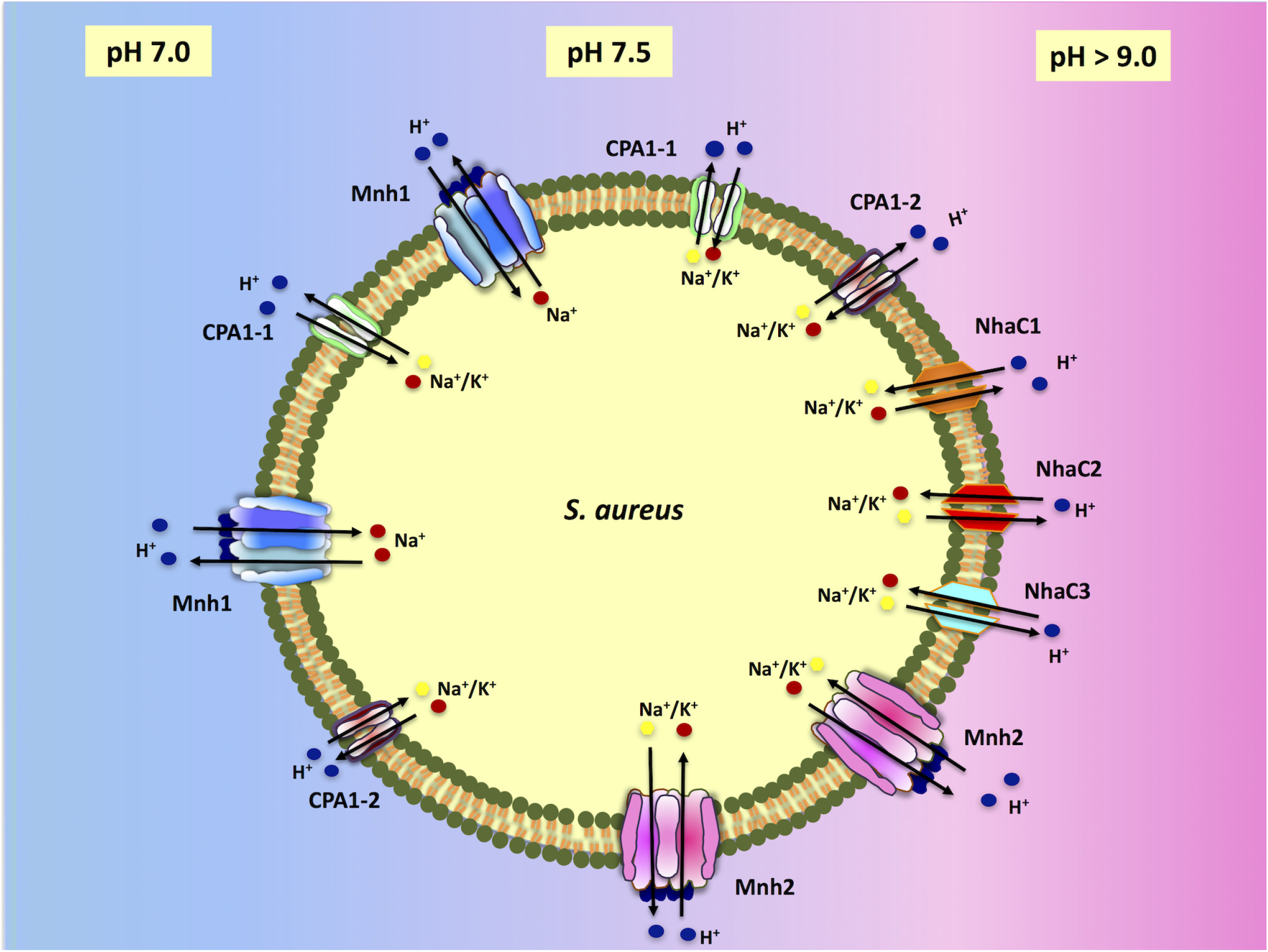
Model depicting whole cohort of secondary cation/proton antiporters in *S. aureus* located at different loci in cytoplasmic membrane. This figure represents two CPA1; CPA1-1 and CPA1-2, two CPA3; Mnh1 and Mnh2 and three NhaCs; NhaC1, NhaC2 and NhaC3 functioning as key players in proton retention in changing environment between pH 6.8 to 9.5 by effluxing Na^+^ and/or K^+^ out in exchange of H^+^. As shown in the figure, Mnh1, CPA1-1 and CPA1-2 actively involved in Na^+^ and/or K^+^ efflux near neutral to pH 7.5. Mnh2, NhaC1, NhaC2, NhaC3 and CPA1-2 exhibited catalytic activity pH > 7.5 to highly alkaline pH.

It has been reported previously that in many other bacterial and archaeal phyla, CPA2 may function as cation/proton efflux system (29). Some of the other CPA2 family transporters have been hypothesized to be channel (30). In *E. coli* and bacillus species CPA2 function is regulated by c-AMP (31). However, functional loss of CPA2 did not affect the viability of *S. aureus* under any pH and salt generated stress conditions (Fig 4A). The data were consistent exhibiting no catalytic activity for K^+^/H^+^ and Na^+^/H^+^ exchange at any pH ranges between pH 7.0 to 9.5 (data not shown). The results strongly suggest that in *S. aureus*, CPA2 does not likely function as cation/proton efflux antiporters but have possibility to function as other transporters or ion channel.

Overall, in *S. aureus* most of the antiporters (CPA1-2, Mnh2, NhaC1, NhaC2 and NhaC3) were regulating pH homeostasis effectively at alkaline pH range. These antiporters were extruding sodium and potassium in exchange of protons to maintain cytoplasmic pH during alkaline stress and sodium toxicity. *S. aureus* is neutralophile and it can successfully survive and grow at higher pH range due to these groups of cation/proton efflux pump that potentially regulates osmotolerance and halotolerance at alkaline pH range (Fig 7). At near neutral pH, Mnh1 was found to synergize with CPA1-1 in order to regulate the sodium toxicity and pH homeostasis while at higher pH range, it was found to functionally synergize with CPA1-2, NhaC1 and NhaC3 that are functionally active at alkaline pH range. The CPA1, CPA3 and NhaC family antiporters have enough functional overlap and each antiporter play critical compensatory role in viability when one has lost function. Inactivation of NhaC3 by clean deletion in Newman significantly reduces the virulence of *S. aureus* in murine infection model. None of other antiporters in NhaC and CPA1 family resulted in marked reduction of virulence. The lethal effect of *nhaC3* was restored by complementing gene in knockout strain. This finding establishes NhaC3 as potential therapeutic targets and provides strong rationale for inhibitor screening against NhaC3 for therapeutic development.

## Acknowledgement

This work was supported by research grant R01GM028454 from the National Institute of General Medical Sciences to T.A.K. A.A. was intern and actively volunteered to this project. Amit Kumar, a junior faculty member at Burke Neurological Institute, contributed in critical proofreading and editing of the manuscript.

